# Representational similarity scores of digits in the sensorimotor cortex are associated with behavioral performance

**DOI:** 10.1101/2021.06.18.448803

**Authors:** J. Gooijers, S. Chalavi, A. Roebroeck, A. Kaas, S.P. Swinnen

## Abstract

Previous studies aimed to unravel a digit-specific somatotopic organization in the primary sensorimotor (SM1) cortex. It is, however, yet to be determined whether such digit somatotopy is associated with motor performance (i.e., effector selection) and digit enslaving (unintentional co-contraction of fingers) during different types of motor tasks. Here, we adopted multivariate representational similarity analysis, applied to high-field (7T) MRI data, to explore digit activation patterns in response to online finger tapping. Sixteen young adults (7 males, mean age: 24.4 years) underwent MRI, and additionally performed an offline choice reaction time task (CRTT) to assess effector selection. During both the finger tapping task (FTT) and the CRTT, force sensor data of all digits were acquired. This allowed us to assess digit enslaving (obtained from CRTT & FTT), as well as digit interference (i.e., erroneous effector selection; obtained from CRTT) and determine the correlation between these variables and digit representational similarity scores of SM1. Digit enslaving during finger tapping was associated with contralateral SM1 representational similarity scores of both hands. During the CRTT, digit enslaving of the right hand only was associated with representational similarity scores of left SM1. Additionally, right hand digit interference was associated with representational similarity scores of left S1. In conclusion, we demonstrate a cortical origin of digit enslaving, and uniquely reveal that effector selection performance is predicted by digit representations in the somatosensory cortex.

## Introduction

To date, there is a vast amount of research supporting that inter-individual differences in brain structure and function are predictive of behavior. Concerning the control of uni- and bimanual movements, previous studies applying neuroimaging techniques, have revealed the involvement of numerous cortical and cerebellar brain regions (for review see Bundy & Leuthardt, 2019; Chettouf, Rueda-Delgado, de Vries, Ritter, & Daffertshofer, 2020; Gooijers & Swinnen, 2014; Swinnen & Wenderoth, 2004). For example, it is generally accepted that hand, and by extension finger, movements elicit increased activation levels in the primary motor cortex (M1) at the level of the hand knob (Yousry et al., 1997). What remains under debate, however, is whether individual finger movements elicit detailed and specific activation patterns within this particular area. While there is consensus in the literature that such a topographical organization of individual digits exists at the level of the post-central primary somatosensory cortex (S1) (Kolasinski et al., 2016; Martuzzi, van der Zwaag, Farthouat, Gruetter, & Blanke, 2014; Sanchez-Panchuelo et al., 2012; Sanchez Panchuelo, Besle, Schluppeck, Humberstone, & Francis, 2018; Schellekens, Petridou, & Ramsey, 2018), evidence for such an organization in pre-central M1 is less explicit. Existing data point towards a) a fine-scale digit somatotopy (Schellekens et al., 2018; Siero et al., 2014) which shows a mirrored pattern of multiple digit representations along the lateral-medial axis at meso-scale (sub-millimeter) resolution (Huber et al., 2020), b) a partly somatotopic arrangement (Beisteiner et al., 2001; Hlustik, Solodkin, Gullapalli, Noll, & Small, 2001; Kleinschmidt, Nitschke, & Frahm, 1997; Lotze et al., 2000), c) a distributed organization with considerable overlap (Indovina & Sanes, 2001; Olman, Pickett, Schallmo, & Kimberley, 2012; Sanes, Donoghue, Thangaraj, Edelman, & Warach, 1995), and d) a complete lack of organization (Schieber, 2002). In sum, the findings on finger representations in M1 are mixed, with the majority reporting evidence of overlapping digit representations that only show a coarse somatotopy when differential contrasts between digits are performed. In order to take into account this distribution of cortical activation patterns of individual digits, a multivariate rather than a univariate approach is better suited to explore cortical representations of digits. Therefore, we investigated digit organization in M1 and S1 using representational similarity analysis (RSA), applied to functional MRI data acquired at high-field strength (7T).

To explore whether these patterns of digit activations at the cortical level are contingent upon everyday hand movements, work by Ejaz and colleagues (2015) revealed that digit representations in M1 are better predicted by natural hand use than by individual musculature. This points towards the fact that overlapping digit patterns at the central level are likely explained by fingers moving together in everyday life to produce natural movements. In addition, the authors reported a moderate negative association between multivariate pattern distances of digit representations in the sensorimotor cortex and digit enslaving (i.e., unintentional co-contraction of non-instructed fingers) when performing an individuated finger press at 75% of maximal voluntary contraction. That is, fingers evoking more similar spatial patterns of activation at the cortical level also showed more co-contraction at the digit level when performing an isometric force task.

Although these studies have significantly improved our insights into the cortical control of finger movements, to our knowledge, no study has attempted to associate digit representations in the sensorimotor cortex with behavioral performance at the level of individual digits. Thus, it is yet to be determined whether cortical representations of individual digits in the sensorimotor cortex are associated with the ability to individually move (i.e., select) an instructed digit and simultaneously actively inhibit neighboring non-instructed digits. Since such tasks rely heavily on effector selection, and planning processes (Van Helvert, Oostwoud-Wijdenes, Geerligs, & Medendorp, 2021), engaging premotor, parietal as well as primary motor cortices (Ariani, Oosterhof, & Lingnau, 2018; Crammond & Kalaska, 2000; Hirose, Nambu, & Naito, 2018; Tanji & Evarts, 1976), we aimed to investigate whether individual digit representations in M1 play a role in effector selection of the contralateral hand. To this end, we included a choice reaction time task (CRTT). Because recent evidence suggests that contralateral S1 is also involved in such movement preparation (Ariani, Pruszynski, & Diedrichsen, 2021; Gale, Flanagan, & Gallivan, 2021), we additionally assessed the importance of digit representations in S1 in effector selection of the contralateral hand. In addition, we determined the association between individual digit representations in the sensorimotor cortex and contralateral digit enslaving. Moving beyond the work by Ejaz and colleagues (2015) in which an isometric force task was used, we aimed to assess the cortical origin of digit enslaving during the performance of a time-pressured CRTT, predominantly dominated by planning processes, as well as during a simple, continuous finger tapping task (FTT), predominantly dominated by pure motor output processes.

We tested the hypothesis that individual digit activation patterns in contralateral M1, and to a lesser extent in contralateral S1, are significantly associated with CRTT performance (reflected as an interference score; i.e., the inability to inhibit movements of non-instructed fingers). More specifically, more overlapping digit activation patterns in the sensorimotor cortex are expected to positively correlate with higher interference scores of the contralateral hand. Additionally, we hypothesized that digit activation patterns in the sensorimotor cortex are associated with digit enslaving (reflected as co-activations of non-instructed fingers) during both the choice reaction time task and the simple, continuous finger tapping task. Finally, whereas previous work generally focused on one hand and one hemisphere, we compared the representational organization of individual digits in the sensorimotor cortex between hemispheres and between regions of interest (M1 and S1), with the expectation to find alike representations in the dominant and non-dominant hemisphere, and more similar activation patterns between digits in M1 relative to S1.

## Results

As the main purpose of our work is to associate individual digit representations in the sensorimotor cortex with behavior at the level of individual digits, we first present the individual activation patterns and representational similarity scores. Next, we discuss the behavioral findings in relation to these representational similarity scores.

### Activity patterns and representational similarity scores

Through analysis of high-resolution functional imaging data, we assessed digit activation patterns in the pre-central and post-central gyri. Figure 1 visualizes activation patterns in response to contralateral finger tapping movements in the left and right hemisphere in a representative participant.

**Figure 1.**
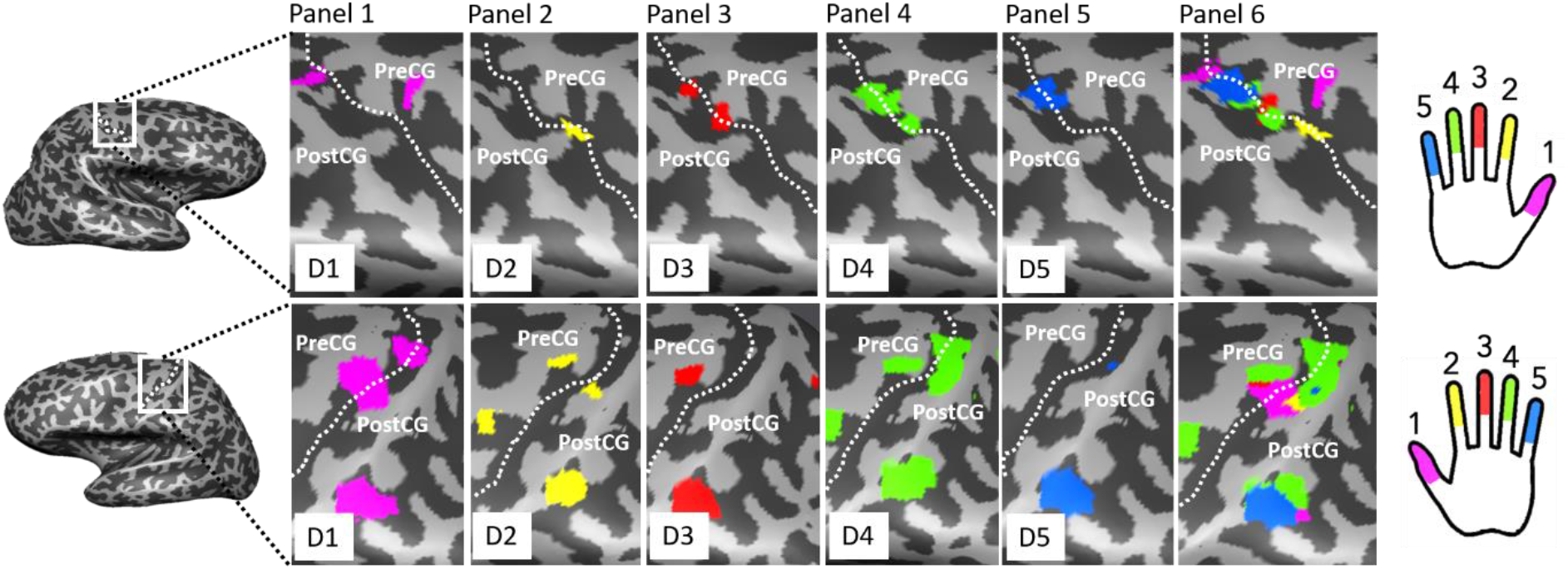
Evoked activation patterns in SM1 during continuous finger tapping with individual fingers for one representative subject. *Left:* Inflated 3D model of right (top) and left (bottom) hemisphere. White square indicates area of pre-central and post-central gyrus. The dotted line indicates the approximate location of the central sulcus. *Middle:* Surface maps for right hemisphere (top) and left hemisphere (bottom) are shown overlaid on the inflated 3D surface model. Panels 1 to 5 display individual digit activation patterns. Panel 6 displays an overlay of all digit activation patterns. The dotted line indicates the approximate location of the central sulcus. Dark grey are areas of negative curvature (sulci), light grey are areas of positive curvature (gyri). Activation patterns represent digit activations contrasted against baseline, with the following thresholds applied (min-max): right hemisphere (5.5-8.0), left hemisphere (3.0-8.0). PreCG = pre-central gryus, PostCG = post-central gyrus. *Right:* a schematic drawing of the color-coded digits of the left (top) and right hand (bottom). Magenta = D1/thumb, Yellow = D2/index finger, Red = D3/middle finger, Green = D4/ring finger, Blue = D5/little finger.

#### Effect of Region of Interest

As assessed by Mantel tests (5000 permutations), similarity matrices of M1 and S1, averaged across participants, were highly related within the left (*r* = .95, *p* = .008) and within the right hemisphere (*r* = .98, *p* = .008). That is, for both hemispheres, the between-digit spatial activation patterns (across all five digits) are highly correlated between M1 and S1. Mantel tests were also performed per participant (left hemisphere: *r* range: .67 to .98; significant (*p* < .05) in 13 out of 15 participants, right hemisphere: *r* range: .09 to .94; significant (*p* < .05) in 14 out of 15 participants). Corresponding correlation plots are presented in the upper panels of Figure 2. Moreover, Wilcoxon Matched Pairs Tests on vectorized correlation coefficients (i.e., vectors of correlation coefficients between digit pairs) revealed higher similarity scores for M1 relative to S1 in the right hemisphere for digit pairs D1-D2, D1-D4, D3-D4, D3-D5, D4-D5 (*ps* < .005), and in the left hemisphere for digit pair D3-D5 (*p* < .005).

**Figure 2.**
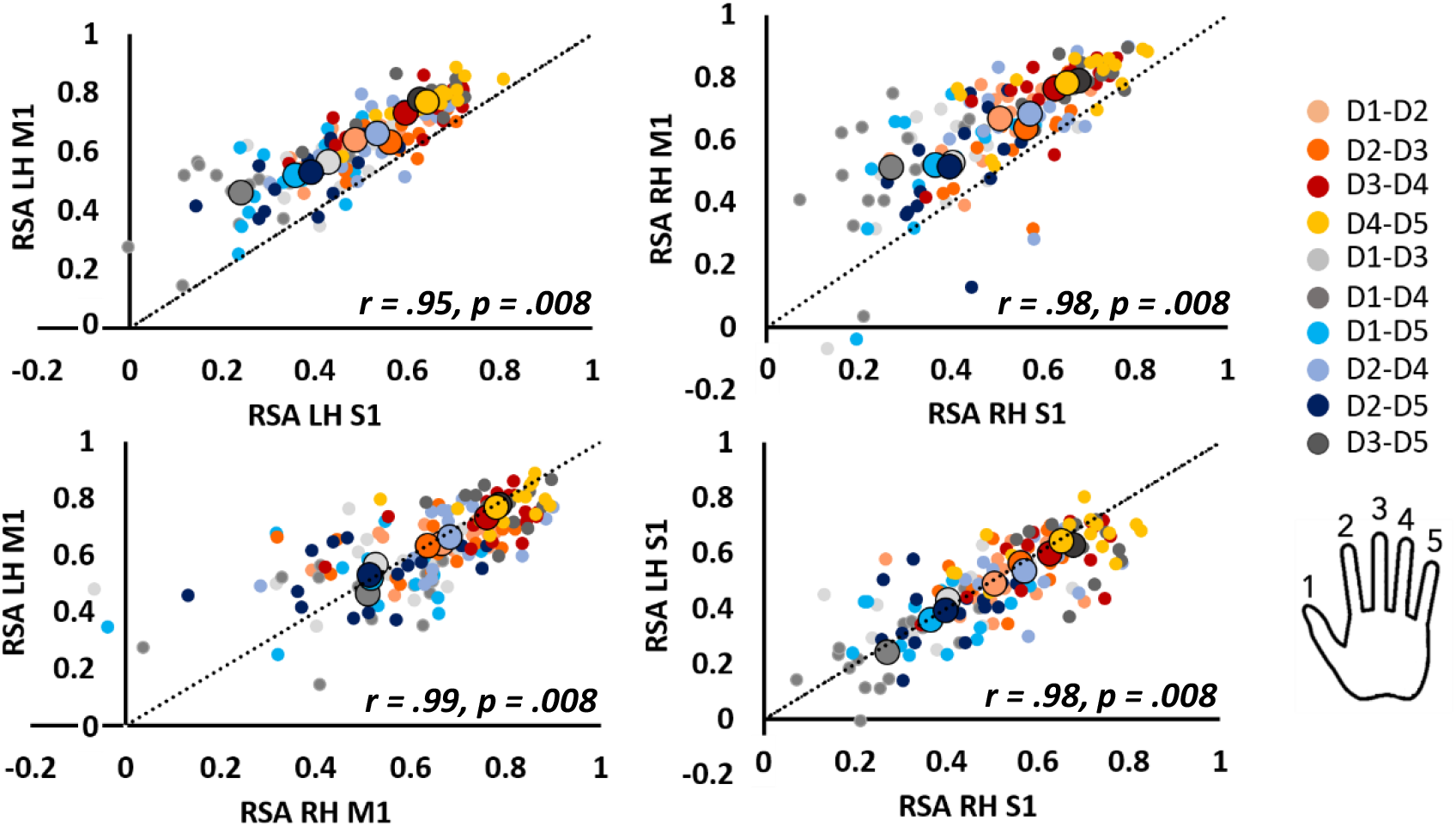
Correlation plots displaying association of RSA values between regions of interest for the left and right hemisphere separately (panels in top row) and between hemispheres for M1 and S1 separately (panels in bottom row). Cool colors refer to non-neighboring digit pairs; warm colors refer to neighboring digit pairs. The dotted line represents an identity line revealing higher RSA values for M1 relative to S1 within hemispheres, but no differences in RSA values between hemispheres within ROIs. Small dots represent individual data points; large dots represent the group average per digit pair. LH = left hemisphere, RH = right hemisphere, M1 = primary motor cortex, S1 = primary somatosensory cortex, RSA = representational similarity analyses. *r* and *p* values represent outcomes of the Mantel Tests across participants.

#### Effect of Hemisphere

As assessed by Mantel tests (5000 permutations), representational structures of the left and right hemisphere, averaged across participants, were highly related for both M1 (*r* = .99, *p* = .008) and S1 (*r* = .98, *p* = .008). Mantel tests were also performed per participant (M1: *r* range: .09 to .96; significant (*p* < .05) in 13 out of 15 participants. S1: *r* range: .12 to .93; significant (*p* < .05) in 12 out of 15 participants). Corresponding correlation plots are presented in the lower panels of Figure 2. Moreover, no hemispherical differences in vectorized correlation coefficients were found for M1 and S1, as assessed by non-parametric Wilcoxon Matched Pairs Tests (M1: *ps* > .05; S1: *ps* > .05).

### Choice Reaction Time Task outside MR scanner – Digit interference and enslaving

Because the digit interference and digit enslaving effects are of interest here, we only report results of these outcome measures. For findings on accuracy and reaction times, we refer the readers to the Supplementary Materials.

#### Digit interference

With regard to vectorized median interference scores, non-parametric Sign Tests revealed no statistically significant difference between Coordination modes (uni- and bimanual; *p* = .61), but revealed that median interference scores of the left hand were significantly higher relative to the right hand (*p* = .010). In addition, a Friedman Test revealed a significant effect of Digit pair, *X*^2^(9) = 95.17, *p* < .001, with the highest interference scores reached for Digit pairs D1-D2, D2-D3, D3-D4, and D4-D5, i.e., the neighboring digits (see Figure 3). In addition, we applied Mantel tests to demonstrate that the interference structure in the left hand was significantly related to the interference structure in the right hand (across participants, *r =* .67, *p* = .042; *r* range: -.33 to .89; significant (*p* < .05) in 3 out of 15 participants). The corresponding correlation plot is presented in Figure 4.

**Figure 3.**
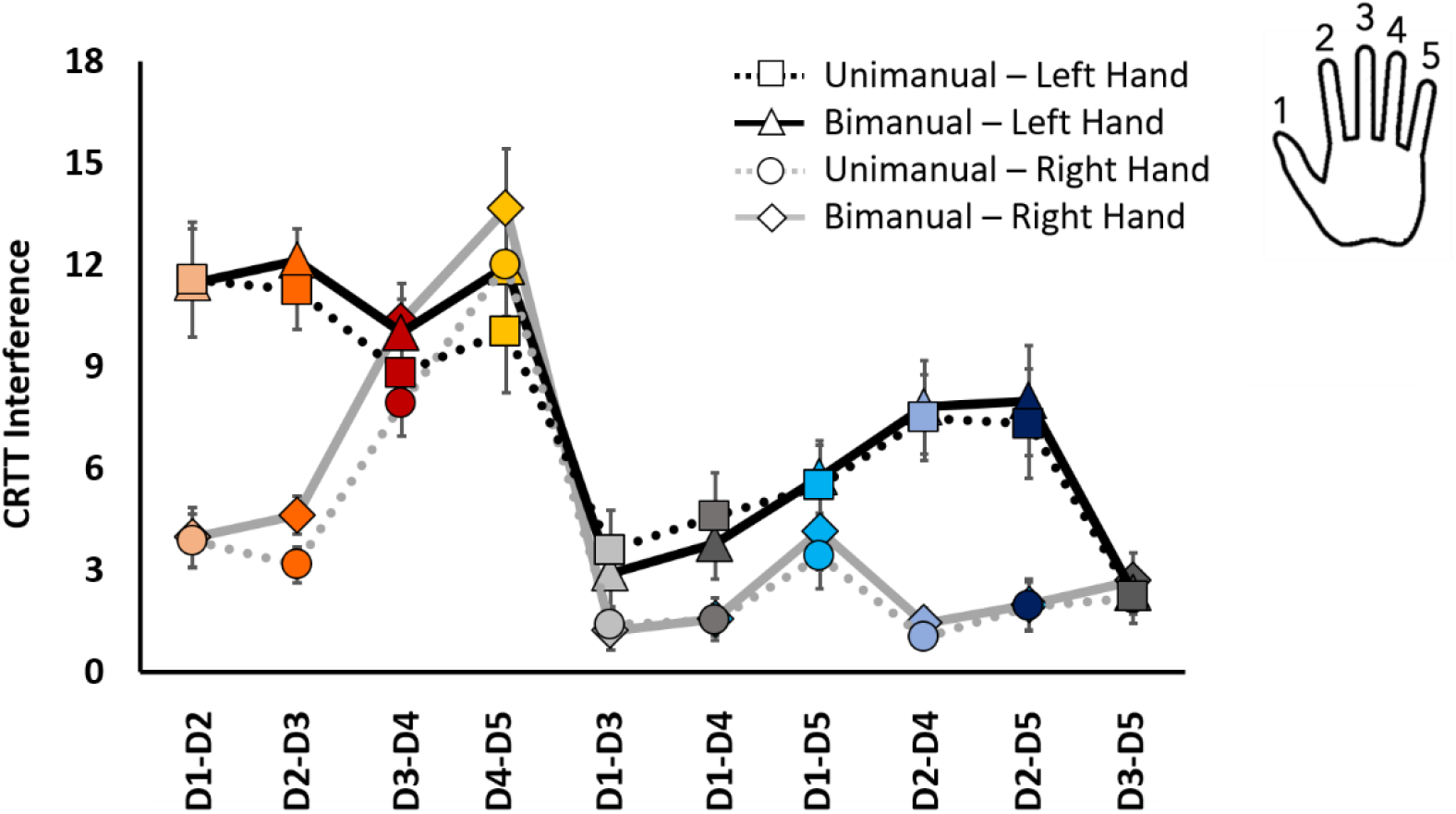
Digit interference scores during CRTT. Error bars indicate standard error of the mean. Cool colors refer to non-neighboring digit pairs; warm colors refer to neighboring digit pairs.

**Figure 4.**
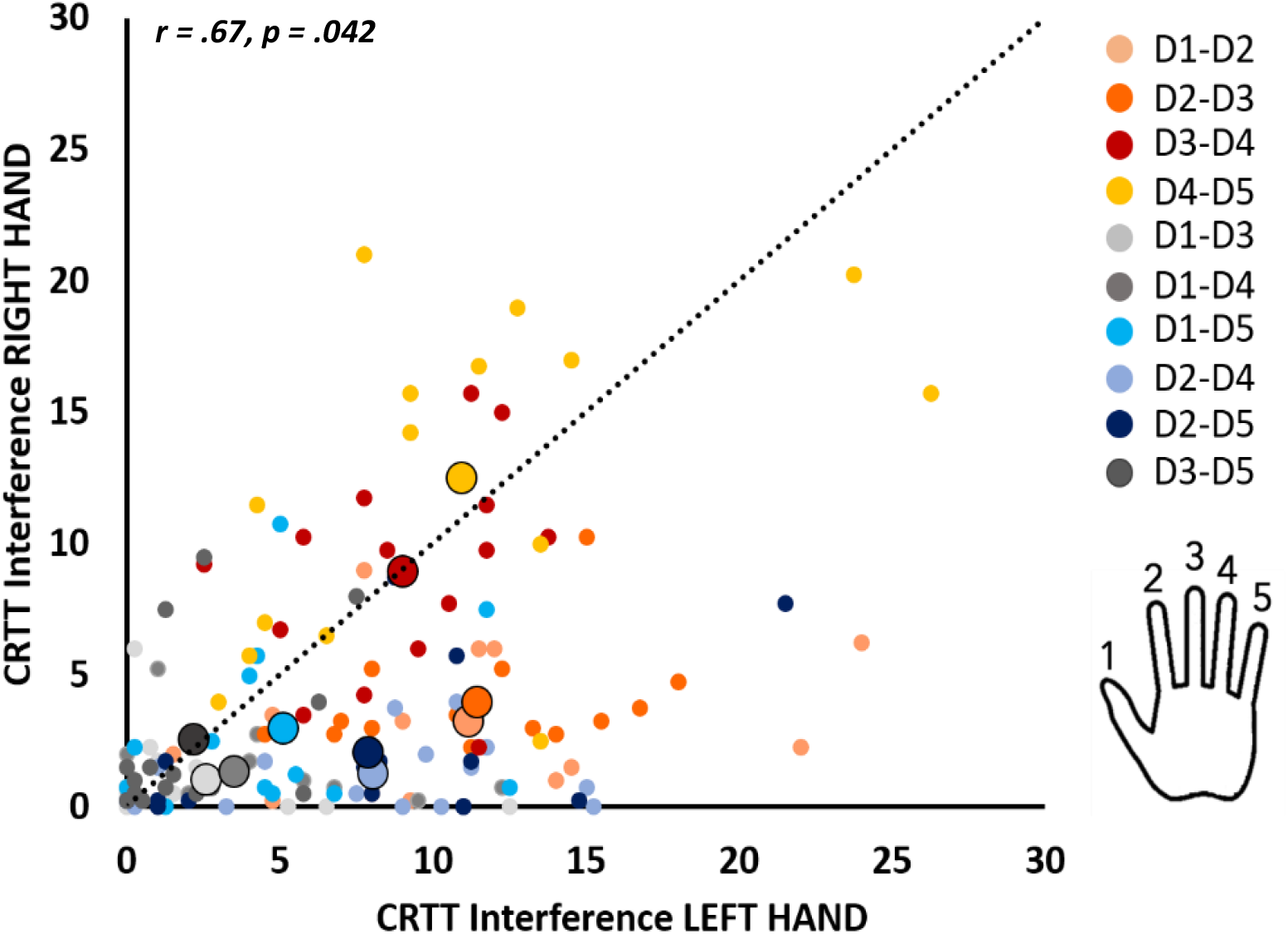
Correlation plot displaying association between CRTT interference scores of the left and right hand. Cool colors refer to non-neighboring digit pairs; warm colors refer to neighboring digit pairs. Small dots represent individual data points; large dots represent the group average per digit pair. The dotted line represents an identity line. With the large colored dots mainly located below the identity line, we demonstrate higher absolute interference scores for the left hand as compared to the right hand. *r* and *p* value represent outcome of the Mantel Test across participants.

#### Digit enslaving

With respect to digit enslaving, non-parametric Sign Tests on vectorized digit pairs revealed no statistically significant differences between Coordination modes (uni- and bimanual; *p* = .61), or between the left and right hand (*p* = .61). However, a Friedman Test revealed a significant effect of Digit pair, *X*^2^(9) = 85.36, *p* < .001, with the highest enslaving scores for digit pairs D3-D4, D3-D5 and D4-D5 (see Figure 5). In addition, we applied Mantel tests to demonstrate that the digit enslaving structure in the left hand was related to the digit enslaving structure in the right hand (across participants, *r* = .85, *p =* .008; *r* range: .15 to .89; significant (*p* < .05) in 6 out of 15 participants). The corresponding correlation plot is presented in Figure 6.

**Figure 5.**
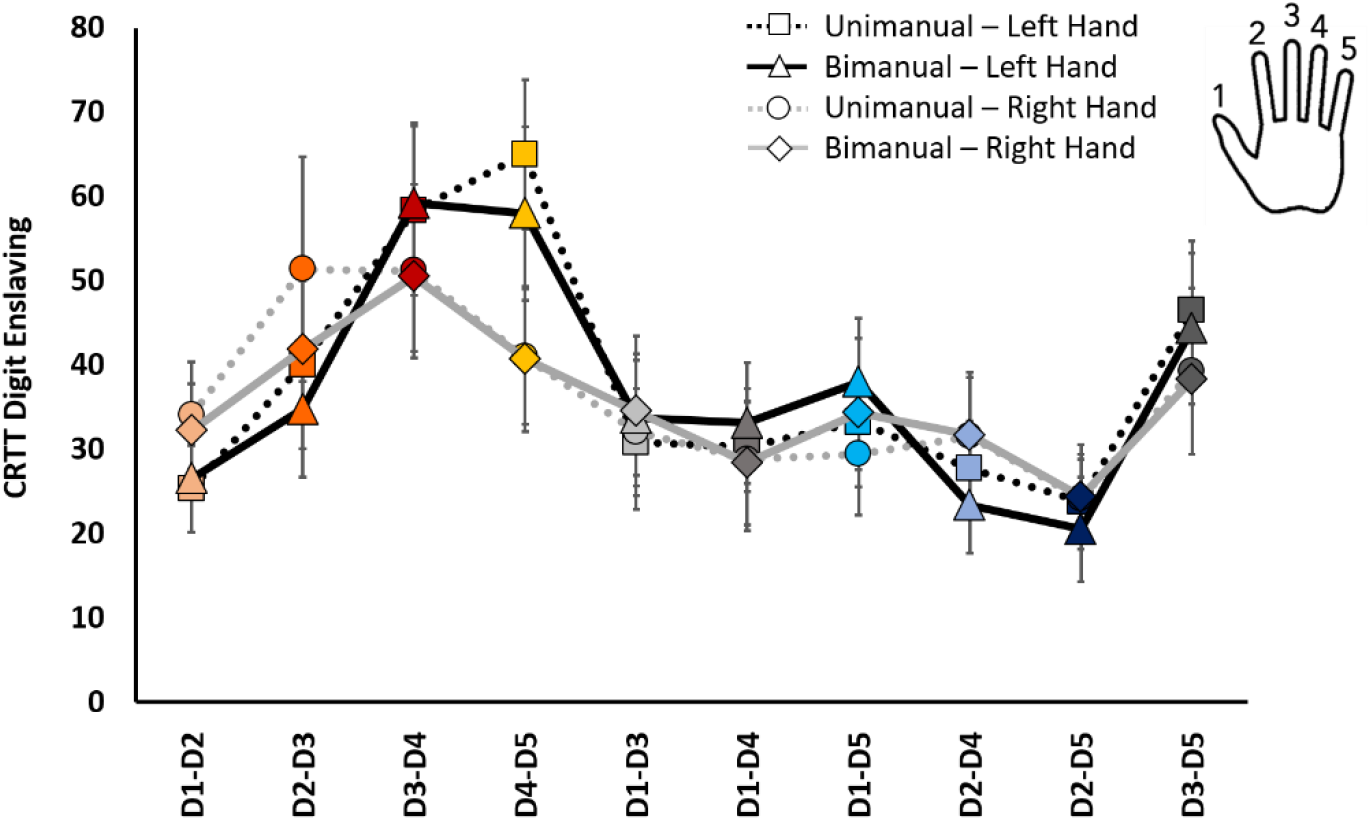
Digit enslaving during CRTT. Error bars indicate standard error of the mean. Cool colors refer to non-neighboring digit pairs; warm colors refer to neighboring digit pairs.

**Figure 6.**
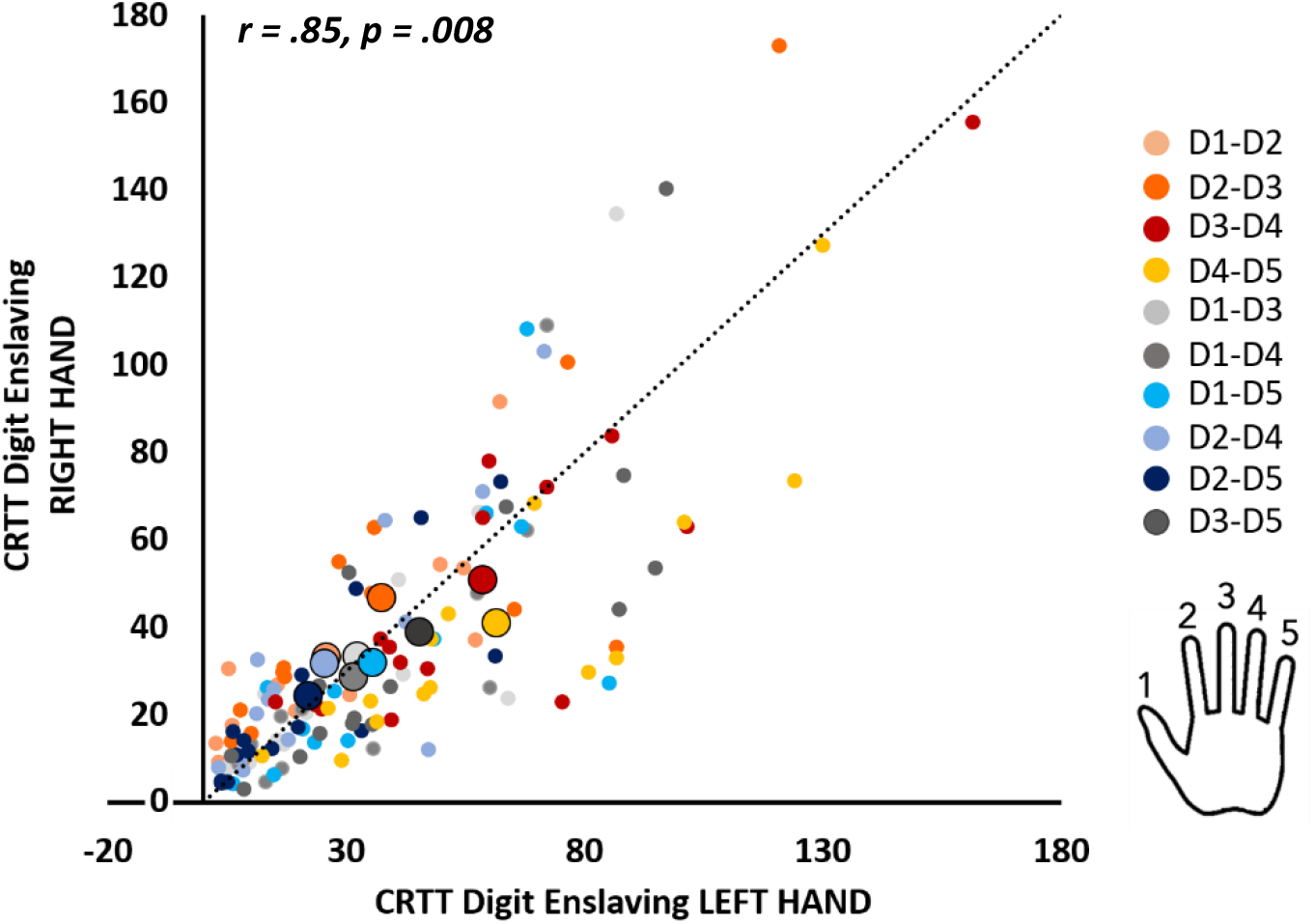
Correlation plot displaying association between digit enslaving of the left and right hand during CRTT. Cool colors refer to non-neighboring digit pairs; warm colors refer to neighboring digit pairs. The dotted line represents an identity line demonstrating no difference in digit enslaving between the two hands. Small dots represent individual data points; large dots represent the group average per digit pair. *r* and *p* value represent outcome of the Mantel Test across participants.

#### Digit Interference and digit similarity scores

As there were no significant differences between interference scores of the uni- and bimanual Coordination modes, we averaged these values and arranged them in a symmetrized matrix for the left and right hand separately. To assess whether interference scores were associated with cortical digit representations, we correlated the CRTT interference matrix with the similarity matrix of the contralateral brain data. Mantel tests with 5000 permutations, averaged across participants, revealed that cortical digit representations of left S1 were significantly correlated with the contralateral CRTT interference matrix; *r* = .59, *p* = .025. This finding indicates that higher similarity scores between activation patterns at the level of S1 (i.e., more overlap between digit representations), are related to more interference (i.e., decreased inhibition of non-cued digits). Other associations did not reach significance; left M1 – right hand: *r* = .44, *p* = .12; right S1 – left hand: *r* = .20, *p* = .28; right M1 – left hand: *r* = .18, *p* = .30. Corresponding correlation matrices and plots are presented in Figures 7A-B, 7G-H and 8, respectively. Mantel tests were also performed per participant (right M1 – left hand: *r* range: *-*.25 to .75; significant (*p* < .05) in 3 out of 15 participants, right S1 – left hand: *r* range: -.25 to .75; significant (*p* < .05) in 3 out of 15 participants, left M1 – right hand: *r* range: *-*.12 to .89; significant (*p* < .05) in 3 out of 15 participants, left S1 – right hand: *r* range: -.24 to .79; significant (*p* < .05) in 5 out of 15 participants).

**Figure 7.**
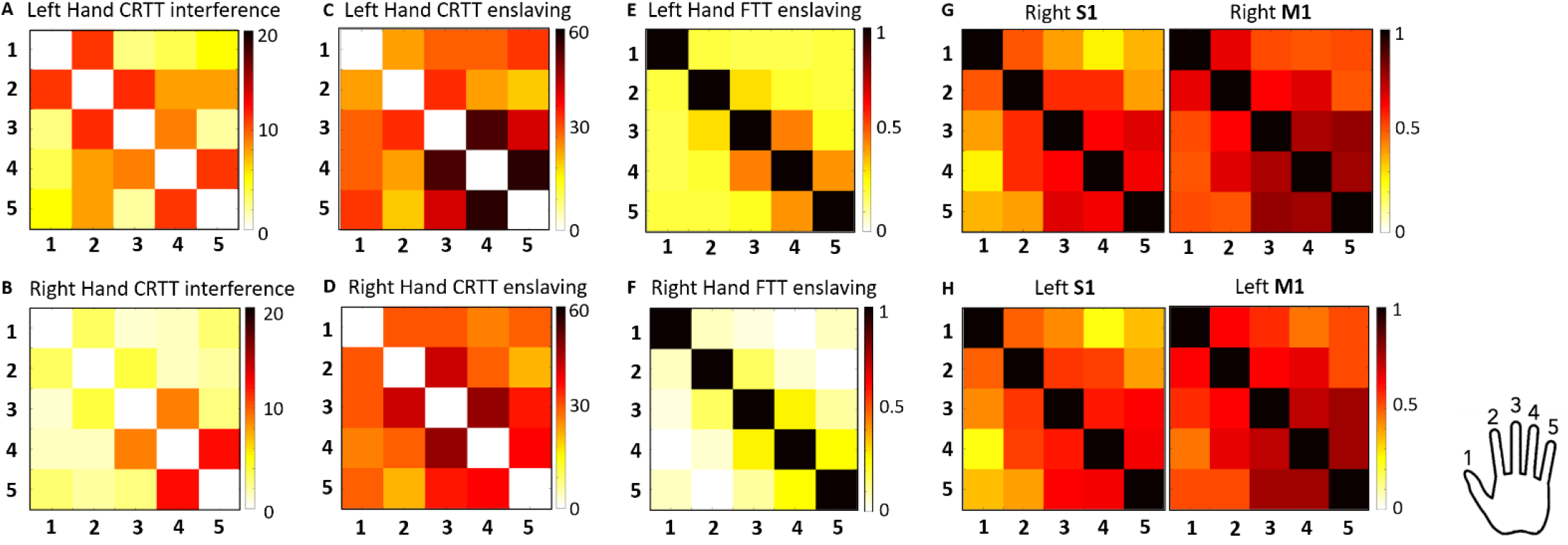
Similarity matrices of group average. **A-B:** Matrices represent interference scores of CRTT arranged into a matrix with darker colors representing more interference between digit pairs. **C-D:** Matrices represent enslaving scores of CRTT arranged into a matrix with darker colors representing more enslaving between digit pairs. **E-F:** Matrices represent enslaving scores of FTT arranged into a matrix with darker colors representing more enslaving between digit pairs. **G-H:** Matrices represent similarity structures of fMRI data for S1 and M1 of the right hemisphere in the top row and S1 and M1 of the left hemisphere in the bottom row. Dark colors represent higher correlation coefficients, i.e., more similar activation patterns between digit pairs. Numbers 1 to 5 represent digits D1 to D5 for rows and columns.

**Figure 8.**
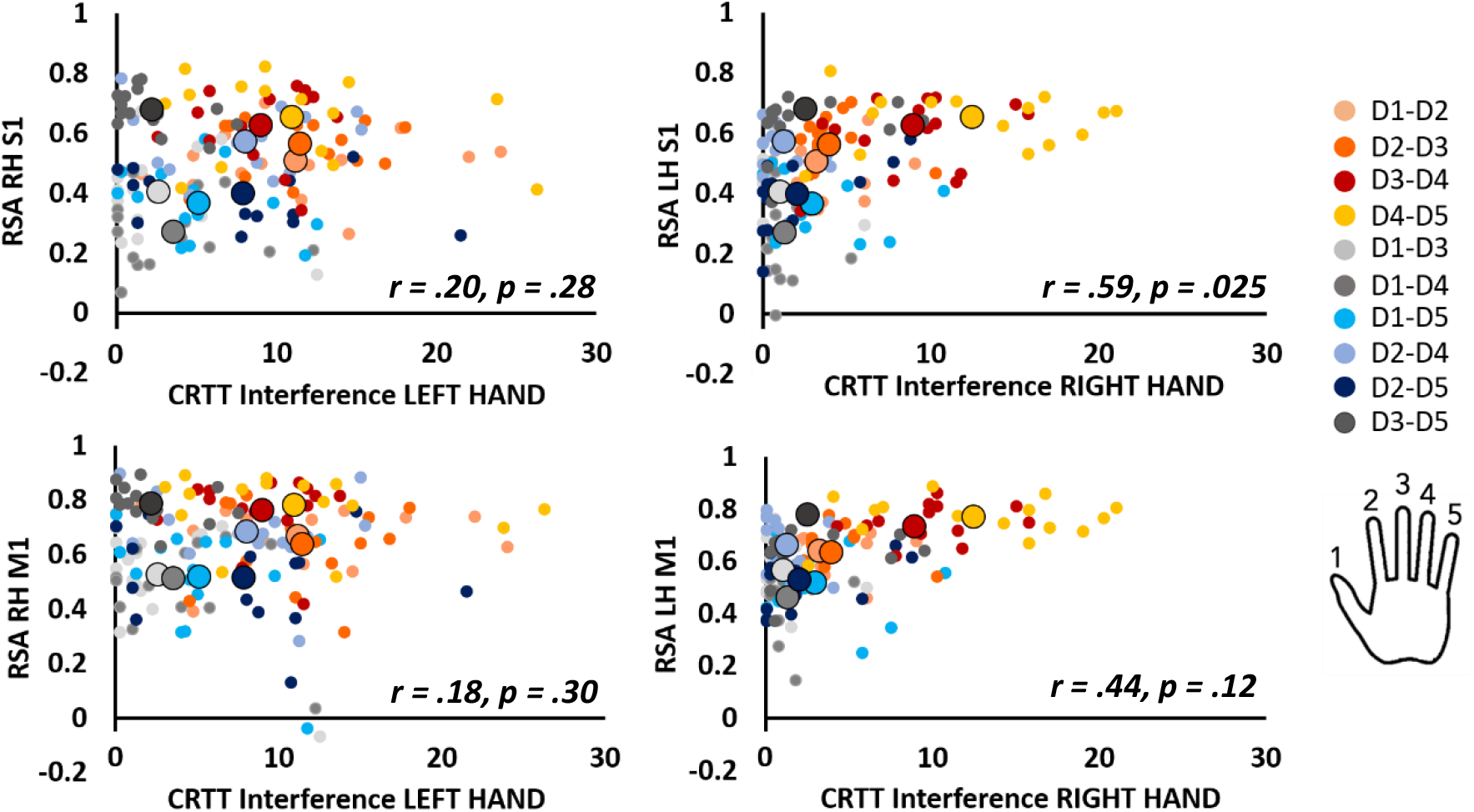
Correlation plots displaying association between digit interference of the left hand (panels on the left) and right hand (panels on the right) during CRTT and RSA values of the contralateral S1 (top row) and M1 (bottom row). Cool colors refer to non-neighboring digit pairs. Warm colors refer to neighboring digit pairs. Small dots represent individual data points; large dots represent the group average per digit pair. RH = right hemisphere, LH = left hemisphere, M1 = primary motor cortex, S1 = primary somatosensory cortex. *r* and *p* values represent outcomes of the Mantel Tests across participants.

#### Digit enslaving and cortical digit similarity scores

As there were no significant differences between digit enslaving for the uni- and bimanual coordination mode, we averaged these values and arranged them in a symmetrized enslaving matrix for the left and right hand separately. To assess whether co-contraction of non-instructed fingers during a choice reaction time task has a cortical origin, we correlated the CRTT digit enslaving matrix with the similarity matrix of the contralateral brain data. Mantel tests with 5000 permutations, averaged across participants, revealed that cortical digit representations of the left hemisphere were predictive of the digit enslaving structure in the contralateral hand; left M1 – right hand: *r* = .64, *p* = .03; left S1 – right hand: *r* = .77, *p* = .02. These findings indicate that higher similarity scores between activation patterns at the level of left M1 and left S1 (i.e., more overlap between digit representations), are associated with more digit enslaving (i.e., co-activation of non-cued digits with cued digit). For the right hemisphere, Mantel tests did not reach the significance level, but demonstrated a similar trend; right M1 – left hand: *r* = .61, *p* = .07; right S1 – left hand: *r* = .60, *p* = .08. Corresponding correlation matrices and plots are presented in Figures 7C-D, 7G-H and 9, respectively. Mantel tests were also performed per participant; right M1 – left hand: *r* range: .08 to .82; significant (*p* < .05) in 5 out of 15 participants, right S1 – left hand: *r* range: -.22 to .77; significant (*p* < .05) in 5 out of 15 participants, left M1 – right hand: *r* range: *-*.01 to .83; significant (*p* < .05) in 4 out of 15 participants, left S1 – right hand: *r* range: -.15 to .85; significant (*p* < .05) in 7 out of 15 participants.

**Figure 9.**
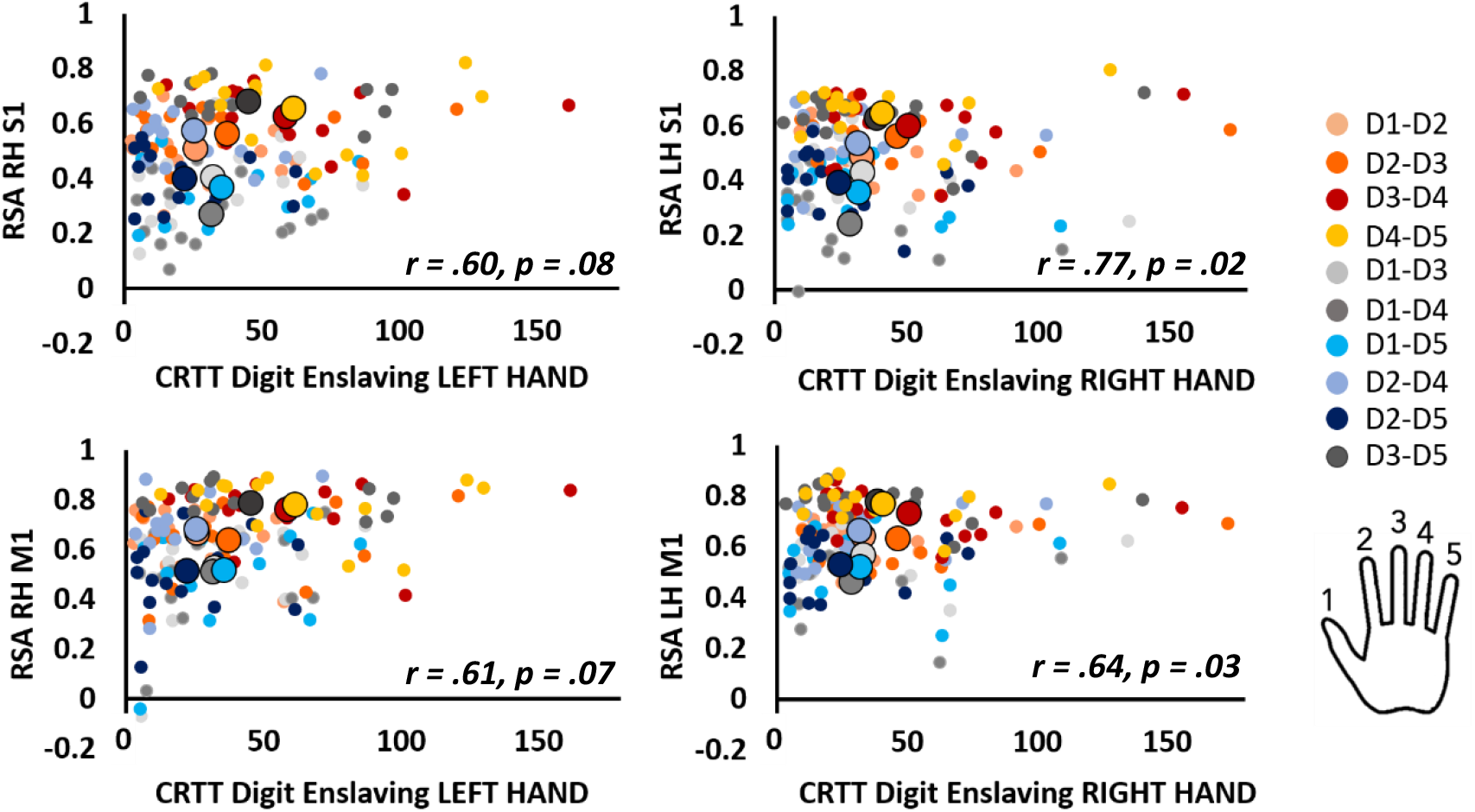
Correlation plots displaying association between digit enslaving of the left (panels on the left) and right (panels on the right) hand during CRTT and RSA values of the contralateral S1 (top row) and M1 (bottom row). Cool colors refer to non-neighboring digit pairs; warm colors refer to neighboring digit pairs. Small dots represent individual data points; large dots represent the group average per digit pair. RH = right hemisphere, LH = left hemisphere, M1 = primary motor cortex, S1 = primary somatosensory cortex. *r* and *p* values represent outcomes of the Mantel Tests across participants.

### Finger Tapping Task inside MR scanner – Digit enslaving

#### Digit enslaving

A repeated-measures ANOVA on the vectorized correlation coefficients between digit pairs during the FTT revealed no main effect of Hand (*F*(1,14) = .04, *p* = .85), a main effect of Digit pair (*F*(9,126) = 26.17, *p* < .001), and no interaction effect between Hand and Digit pair (*F*(9,126) = .86, *p* = .56). Tukey post-hoc pairwise comparisons revealed that digit pairs D2-D3, D3-D4 and D4-D5 reached higher correlation coefficients relative to all other digit pairs (*ps* < .001). Results are presented in Figure 10. In addition, we used Mantel tests to demonstrate that the structure of FTT digit enslaving in the left hand was related to the structure of FTT digit enslaving in the right hand (across participants, *r =* .85, *p* = .008; *r* range: .16 to .99; significant (*p* < .05) in 9 out of 15 participants). The corresponding correlation plot is presented in Figure 11.

**Figure 10.**
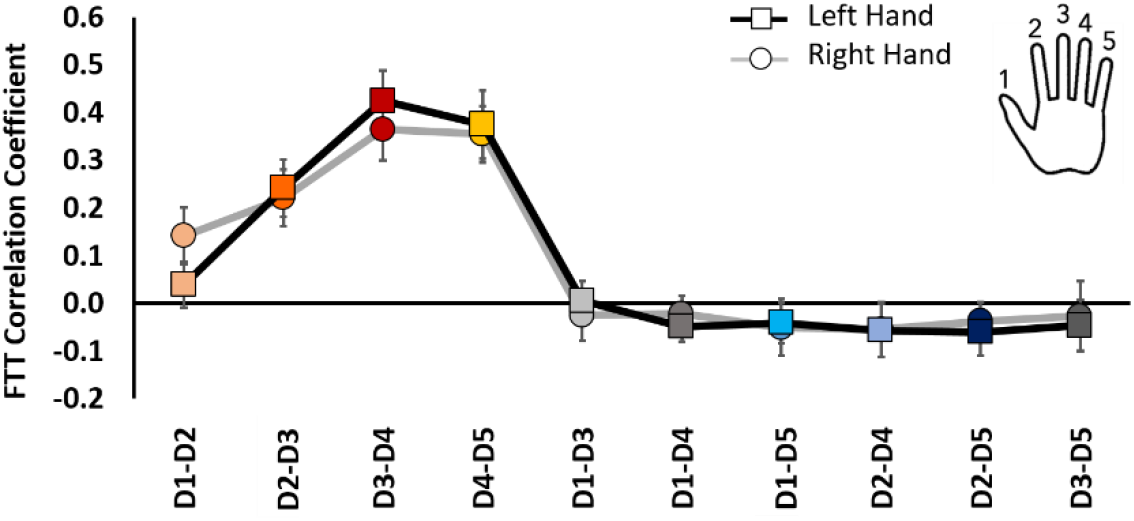
*Digit enslaving during* FTT. Error bars indicate standard error of the mean. Cool colors refer to non-neighboring digit pairs; warm colors refer to neighboring digit pairs.

**Figure 11.**
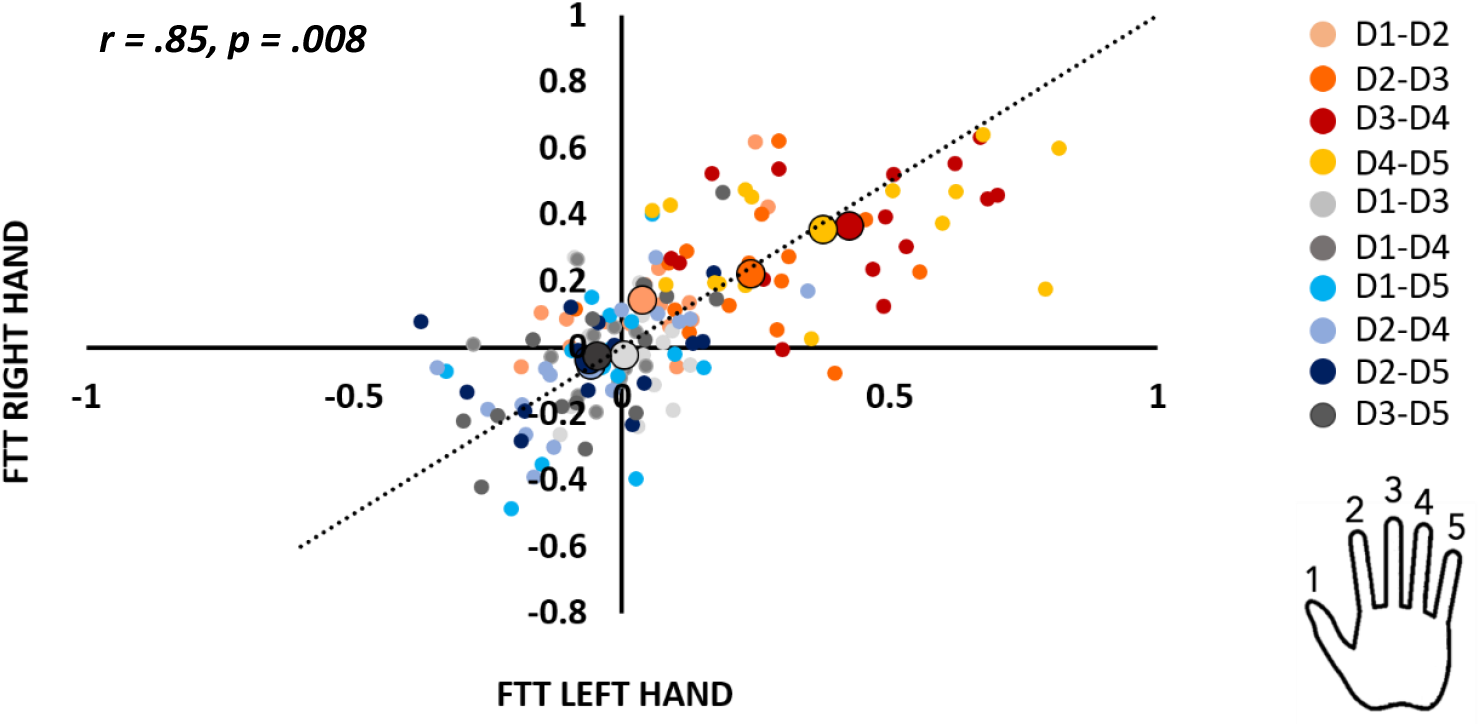
Correlation plot displaying association between digit enslaving of the left and right hand during FTT. Cool colors refer to non-neighboring digit pairs. Warm colors refer to neighboring digit pairs. The dotted line represents an identity line demonstrating no difference in digit enslaving between the two hands. Small dots represent individual data points; large dots represent the group average per digit pair. *r* and *p* value represent outcome of the Mantel Test across participants.

#### Digit enslaving and cortical similarity scores

To assess whether co-contraction of non-instructed fingers during a continuous finger tapping task has a cortical origin, we correlated the FTT enslaving matrix with the similarity matrix of the contralateral brain data. Mantel tests with 5000 permutations, averaged across participants, revealed that cortical representations were correlated with the FTT digit enslaving pattern of the contralateral hand; right M1 – left hand: *r* = .78, *p* = .03; right S1 – left hand: *r* = .81, *p* = .03; left M1 – right hand: *r* = .75, *p* = .03; left S1 – right hand: *r* = .84, *p* = .01. These findings indicate that higher similarity scores between activation patterns at the level of the sensorimotor cortex (i.e., more overlap between digit representations), are associated with more contralateral digit enslaving (i.e., co-activation of non-cued digits with cued digit). Correlation matrices and plots are presented in Figures 7E-F, 7G-H and 12, respectively. Mantel tests were also performed per participant (right M1 – left hand: *r* range: *-*.02 to .84; significant (*p* < .05) in 7 out of 15 participants; right S1 – left hand: *r* range: -.04 to .88; significant (*p* < .05) in 4 out of 15 participants, left M1 – right hand: *r* range: .-.05 to .82; significant (*p* < .05) in 5 out of 15 participants, left S1 – right hand: *r* range: .09 to .90; significant (*p* < .05) in 7 out of 15 participants).

**Figure 12.**
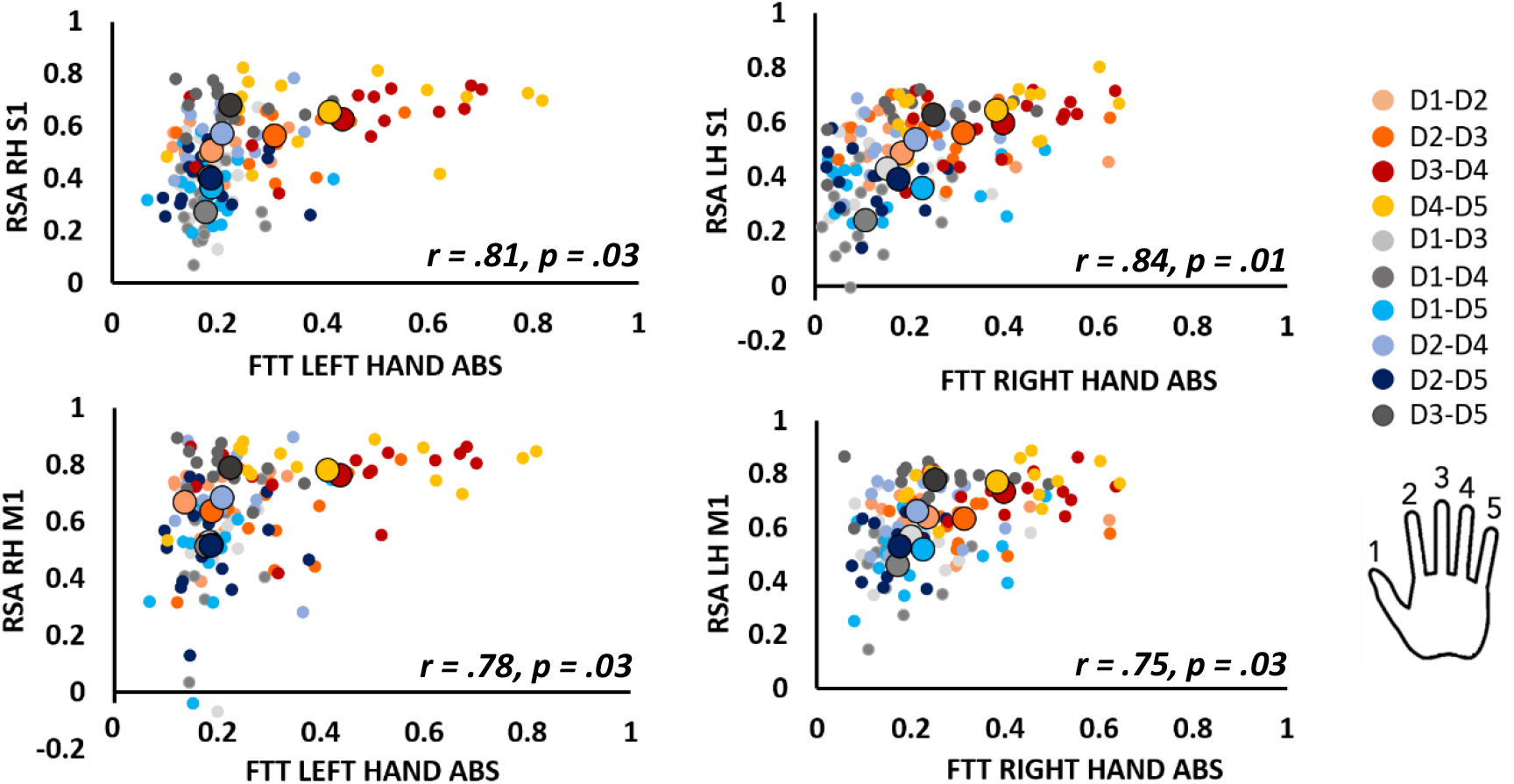
Correlation plots displaying association between digit enslaving of the left (panels on the left) and right (panels on the right) hand during FTT and RSA values of the contralateral S1 (top row) and M1 (bottom row). Cool colors refer to non-neighboring digit pairs. Warm colors refer to neighboring digit pairs. Small dots represent individual data points; large dots represent the group average per digit pair. RH = right hemisphere, LH = left hemisphere, M1 = primary motor cortex, S1 = primary somatosensory cortex, ABS = absolute correlation coefficients. *r* and *p* values represent outcomes of the Mantel Tests across participants.

## Discussion

Here, we applied a multivariate approach to explore cortical representations of digits in the pre-central and post-central gyri and investigated how these relate to specific motor behavior at the level of individual digits. We found that digit enslaving during continuous finger tapping was significantly associated with contralateral SM1 representational similarity scores of both hands. During the choice reaction time task, digit enslaving of the right hand only was significantly associated with representational similarity scores of the contralateral left SM1. Additionally, right hand digit interference (i.e., inability to inhibit non-cued fingers) was significantly associated with representational similarity scores of left S1. In conclusion, we demonstrate a cortical origin of digit enslaving and uniquely reveal that effector selection (i.e., planning) during CRTT (as reflected by digit interference) is predicted by digit representations in the contralateral somatosensory cortex.

### Behavior

With the *digit interference* score, obtained from the CRTT, we aimed for a measure that indicates the failure to preserve selective motor inhibition (Prigatano et al., 2020), reflecting the preparatory process of effector selection. Unlike digit enslaving levels during the performance of the CRTT (see further), digit interference revealed a significant difference between the dominant and non-dominant hand, with a reduction in adequately inhibiting inappropriate (non-cued) responses for digits of the non-dominant hand. Although these findings are in contrast with recent findings by Prigatano et al., (2020) who applied the Halstead Finger Tapping Test, our findings are consistent with the phenomenon of surround inhibition (Beck & Hallett, 2011) being more pronounced in the dominant hemisphere of right-handed participants. That is, a more efficient selective execution of desired movements is achieved in the dominant hand (Shin, Sohn, & Hallett, 2009). Although one could argue a potential floor effect in the dominant right hand, particularly for neighboring digits there seems to be room for improvement concerning the interference scores. Despite that absolute interference scores showed a marked difference between hands, we found a highly comparable overall pattern of interference scores across digits between the left and right hand. Not surprisingly, neighboring digits revealed higher interference scores relative to non-neighboring digits across both hands.

During both the time-pressured CRTT and the continuous FTT, we demonstrated that the highest *digit enslaving* (i.e., co-activation) levels are particularly present for neighboring digits including D2, D3, D4 and D5. D1 (i.e., thumb) represents a rather distinct pattern in comparison to all other digits, suggesting rather independent behavior. These findings extend the work by Ejaz and colleagues (2015) which similarly revealed enslaving of adjacent fingers, albeit when performing a single finger maximal voluntary contraction. These results indicate that, irrespective of task variations such as finger lifting versus finger pressing, co-movements of adjacent fingers, in particular, are present. Noteworthy, we did not observe significant differences in digit enslaving between the non-dominant and dominant hand. This is in line with the majority of previous studies reporting no (Hager-Ross & Schieber, 2000; Kimura & Vanderwolf, 1970; Reilly & Hammond, 2004; Wilhelm, Martin, Latash, & Zatsiorsky, 2014) or very small differences in digit enslaving between the two hands (Li, Danion, Latash, Li, & Zatsiorsky, 2000a, 2000b). The absence of a significant difference in digit enslaving between the dominant and non-dominant hand suggests that the more extensive and more fine-grained use of the dominant hand in everyday motor activities relative to the non-dominant hand, does not result in higher levels of digit enslaving (Wilhelm et al., 2014). Of note, digit enslaving in the CRTT was measured in a sitting position, whereas digit enslaving in the FTT was measured in a supine position (in the MR scanner). As the enslaving matrices resulting from CRTT and FTT performance were highly correlated, this suggests that there is little to no effect of participant posture and task paradigm on digit enslaving at the behavioral level.

### Cortical activation patterns

We observed that the representations of activity patterns in M1 and S1 in response to individuated finger movements were markedly similar. That is, the relative distribution of brain activity representing individual digit movements within one hand is comparable between M1 and S1 within the same hemisphere. Notably, within the non-dominant right hemisphere and particularly for neighboring digit pairs, absolute correlation coefficients were significantly higher in M1 relative to S1. A similar trend was observed in the dominant left hemisphere, although not surviving corrections for multiple testing. These findings indicate that although the pattern was comparable, the activity patterns of single digits revealed higher similarity scores in M1 than in S1. This was expected based on previous studies reporting a more distinct homunculus-like pattern of digit activations in the post-central relative to the pre-central gyrus (Huber et al., 2020; Martuzzi et al., 2014; Sanchez Panchuelo et al., 2018). This is likely the result of everyday coordinated use of digits when for example grasping, writing, or picking up objects. The hemispheric difference, i.e., a significant difference between M1 and S1 in the right, but not the left hemisphere is likely the result of the right hemisphere mainly controlling the non-dominant left hand, which may predispose simultaneous over individuated finger movements given its stabilizing, supporting role in many coordinated movements requiring division of labor between hands. As a result hereof, more overlap between digit representations is anticipated in M1, prompting a larger difference with S1 in which individual representations of digits are clearly distinguishable irrespective of everyday movements.

Importantly, our findings did not reveal a significant lateralization effect when directly comparing digit representations in M1 left with M1 right, and in S1 left with S1 right. This suggests that it is particularly the within-hemisphere interplay between M1 and S1, structurally connected through U-shaped white matter fibers (Catani et al., 2012; Guevara et al., 2011; Shinoura et al., 2005), that is subject to experience-induced changes. As similarity scores are hardly studied in both hemispheres, our data add important new knowledge indicating that although the representational patterns were highly correlated between M1 and S1 in both hemispheres, particularly in the non-dominant right hemisphere higher correlation values between digits (suggesting more overlap) were found in M1 compared with S1. Of note, unlike the majority of the work on digit somatotopy in S1, we assessed digit representations based on active finger movements, and not passive stroking of digits.

### Association behavior and cortical activation patterns

#### The cortical origin of digit enslaving

To the best of our knowledge, our findings demonstrate for the first time that the digit enslaving matrix of the right hand during the performance of a time-pressured CRTT was significantly associated with the pattern of digit representations in the contralateral sensorimotor cortex. This finding significantly adds to work by Ejaz and colleagues (2015), revealing a neural source of digit enslaving while performing a maximal isometric force production task. Although correlations between digit enslaving of the left hand and digit activation patterns of right M1 and S1 were trending towards significance as well, there is a potential influence of body side/hemisphere. As attested by the correlation matrices and plots, it is likely that the source of this hemispheric difference is found at the behavioral level.

Moreover, further supporting the cortical origin of digit enslaving, we reported high correlations between FTT digit enslaving of the left and right hand and representational similarity matrices of the contralateral sensorimotor cortices. Although it is to some extent surprising that digit enslaving for FTT, but not for CRTT, revealed strong associations with neural representations in both hemispheres, it should be noted that the representational similarity analyses were performed on data acquired during FTT and not CRTT performance (which was performed outside the MR scanner). Moreover, whereas the FTT requires repetitive tapping and is therefore easy to sustain and less error-prone, the CRTT does instigate errors as participants are forced to make a choice under time pressure. Although we merely included trials in which the cued finger was correctly lifted, digit enslaving values could have been influenced by the erroneous lifting of neighboring digits.

Although the cortical origin of digit enslaving was previously suggested (Lang & Schieber, 2004; Yu, van Duinen, & Gandevia, 2010; Zatsiorsky, Li, & Latash, 2000), the first evidence was published only recently and demonstrated stronger digit enslaving to be related with more similar activation patterns in the sensorimotor cortex, by employing multi-voxel pattern fMRI analyses (Ejaz et al., 2015). However, digit enslaving in the latter study was assessed during isometric force production with right hand digits only in a small sample (n=7). Here, we move beyond their findings by uniquely demonstrating that more similar activation patterns between digits in the sensorimotor cortices of both hemispheres are predictive of higher digit enslaving in the contralateral hands during a continuous finger tapping and a time-pressured choice reaction time task.

#### Digit interference is partly predicted by digit activation patterns in the sensorimotor cortex

It was anticipated that the ability to independently move a cued finger, and at the same time inhibit non-cued fingers (reflecting effector selection processes), would be associated with the cortical digit activation patterns of the contralateral primary sensory and particularly motor cortex. That is, the more similar activation patterns for digit pairs are, the more likely the occurrence of incorrect (co-)lifting of non-cued fingers. In the present dataset, a significant positive association was found between the similarity scores of left S1 and interference scores of the right (dominant) hand digits. This suggests that higher digit interference (i.e., reduced inhibition of (adjacent) non-cued fingers) was significantly associated with more overlapping activation patterns in S1 in particular. Although a more prominent association was principally anticipated in M1, it is not surprising that the somatosensory cortex is involved as participants were asked to fixate on the screen and not look at their fingers, forcing them to rely on proprioceptive and tactile feedback to perform the CRTT. Furthermore, recent work provides evidence that also S1 changes its neural state to prepare for proprioceptive and sensory signals to occur as a result of movement execution (Gale et al., 2021) and to predict the planned finger movement (Ariani et al., 2021). Nonetheless, the concept of surround inhibition, a process likely occurring during the motor planning phase (Beck & Hallett, 2010) is potentially more dependent on higher-order motor areas and/or fronto-thalamic pathways than on digit similarity scores in SM1. Finally, interference scores were possibly impacted by visual discrimination errors (i.e, incorrectly differentiating between neighboring cued digits on the screen). However, relatively large interference scores were also observed for the most peripheral digits (D1-D2 & D4-D5), which should be visually discriminated more easily relative to digits D2, D3 and D5. Thus, we may infer from our findings that for CRTT digit interference, contralateral representational digit activation patterns are particularly relevant in the sensory cortex and, unlike for digit enslaving, less so in the motor cortex. Potentially, visual processing or higher-order motor areas are more involved in this process of digit interference. Future studies are, however, necessary to confirm these indications.

As a final remark, we would like to point towards the substantial inter-individual variability that was observed for the majority of Mantel outcomes (Mantel tests were used to assess the association between two correlation matrices at the neural and the behavioral level). This suggests that despite including a fairly homogeneous sample, unique individual experiences appear to shape digit patterns at the behavioral as well as the neural level, while retaining general similarity scores at the group level.

## Conclusion

Overall, our findings suggest a mapping between topographical digit representations in SM1 and contralateral digit enslaving behavior. Additionally, we provide a first indication in a healthy young population that the ability to individually move a cued digit and simultaneously actively inhibit non-cued digits (i.e., effector selection/motor planning) is associated with the amount of overlap in the distributed mapping of these individual digits in S1, but not M1.

## Materials and methods

### Participants

A total of 16 healthy young adults were included in the present study (7 males, mean age = 24.4 years, SD = 2.7, range = 21.4 – 30.3). All participants were right-handed according to the Edinburgh Handedness Scale (Oldfield, 1971). The mean laterality quotient was 90.6% (SD = 12.9, range 70-100). Participants were excluded from the study in case of usage of psychoactive medication, a history of drug abuse, psychiatric or neurological disease, significant multiple trauma, or in the case of MRI contraindications. Additionally, given the potential effect of years of musical training on finger representations in the sensorimotor cortex, musical experience was assessed by means of the Musical Experience Questionnaire (J. Bailey & Penhune, 2012; J. A. Bailey & Penhune, 2010). Participants who reported playing a musical instrument or had experienced more than 3 years of formal musical training at the time of testing, were excluded. All participants were informed about the study and provided written informed consent prior to participation, according to the Declaration of Helsinki. All experimental procedures were approved by the ethics committee in Leuven (S58357) and the Medical Ethical Committee of the Maastricht University.

### Experimental design

Two sessions were administered: a behavioral session which took place at the Motor Control Laboratory of the KU Leuven, and an imaging session at the Maastricht Brain Imaging Center, within an average time window of 9 days (SD = 5.61).

#### Behavioral procedure

In our experiment, two behavioral tasks were included, i.e., the Choice Reaction Time Task (CRTT) and the Finger Tapping Task (FTT). These tasks were selected as they are complementary in terms of underlying processes during effector movements. Whereas the first relies profoundly on effector selection processing (i.e., planning), the latter is mainly involved with effector output processes (i.e., execution). Below, we provide more detail for each of these tasks.

### Choice Reaction Time Task

In order to assess the association between individual digit representations in the sensorimotor cortex and the ability to accurately and efficiently select (i.e., plan) and move individual cued digits, we included a Choice Reaction Time Task (CRTT) in our experimental design. More specifically, participants were asked to lift a particular digit as fast as possible in response to a specific visual stimulus. During CRTT performance, participants were seated comfortably at a table in front of a PC screen with their lower arms positioned on the table and fingertips resting on a set of force transducers. A custom-made set-up consisting of ten force transducers (Honeywell FS series, (based on Ejaz et al., 2015)) pinned on a plate (Figure 13A), was positioned on the table in front of the participant. For each force transducer, a 3D-printed acrylonitrile Butadiene Styrene holder with a peg at the bottom was created, to fixate the transducers to the plate. This allowed for adjustment of the force transducers to ensure a natural position of the fingers for each participant and easy handling of the force sensors. During the task, participants were presented with multiple stimuli (i.e., ten individual digits) on the monitor in front of them and were required to respond as fast and as accurately as possible to the cued stimulus/digit with the corresponding finger. More specifically, at the start of each run, a message was presented on the monitor in front of the participant (duration of 2 seconds) warning the participant to prepare for the upcoming trials. Next, two schematic white hands with a fixation cross in the middle were displayed. Participants were instructed to fixate on the cross throughout the task. At the start of each trial, one finger switched to a red color (= visual cue), indicating to the participant to lift the respective finger from the force transducer, and to smoothly return the responding finger to its neutral position afterward. After each trial, there was a short delay of 1.33 seconds (movement frequency was 0.75 Hz), before the next cue was presented. The order of visual cues was pseudo-randomized. The CRTT was performed in 3 runs; (#1) unimanual left hand, (#2) unimanual right hand, and (#3) bimanual. For the unimanual runs (#1 and #2), 150 trials were included, presented to the participant in three blocks of 50 trials, with a rest period of 18 seconds in between. For the bimanual run (#3), 300 trials were included, presented to the participants in six blocks of 50 trials, with a rest period of 18 seconds in between. In total, participants responded 90 times with each finger. An overview of the CRTT task conditions is provided in Figure 13B. Participants were allowed a short practice run to familiarize themselves with the set-up and task. In order to calibrate the force transducers, baseline measurements (i.e., no fingers on the force transducers) were taken prior to the first run, and after the last run. As all force transducers were set up to continuously register force levels throughout the task, and participants were instructed to keep all fingers on the force transducers at all times (except when cued), co-activation of non-cued fingers was also measured. This experimental set-up allowed us to assess digit enslaving (i.e., the degree to which a cued digit is able to move individually while non-cued fingers remain stationary) and digit interference (see “Analysis of behavioral data”) during a time-pressured reaction time task.

**Figure 13.**
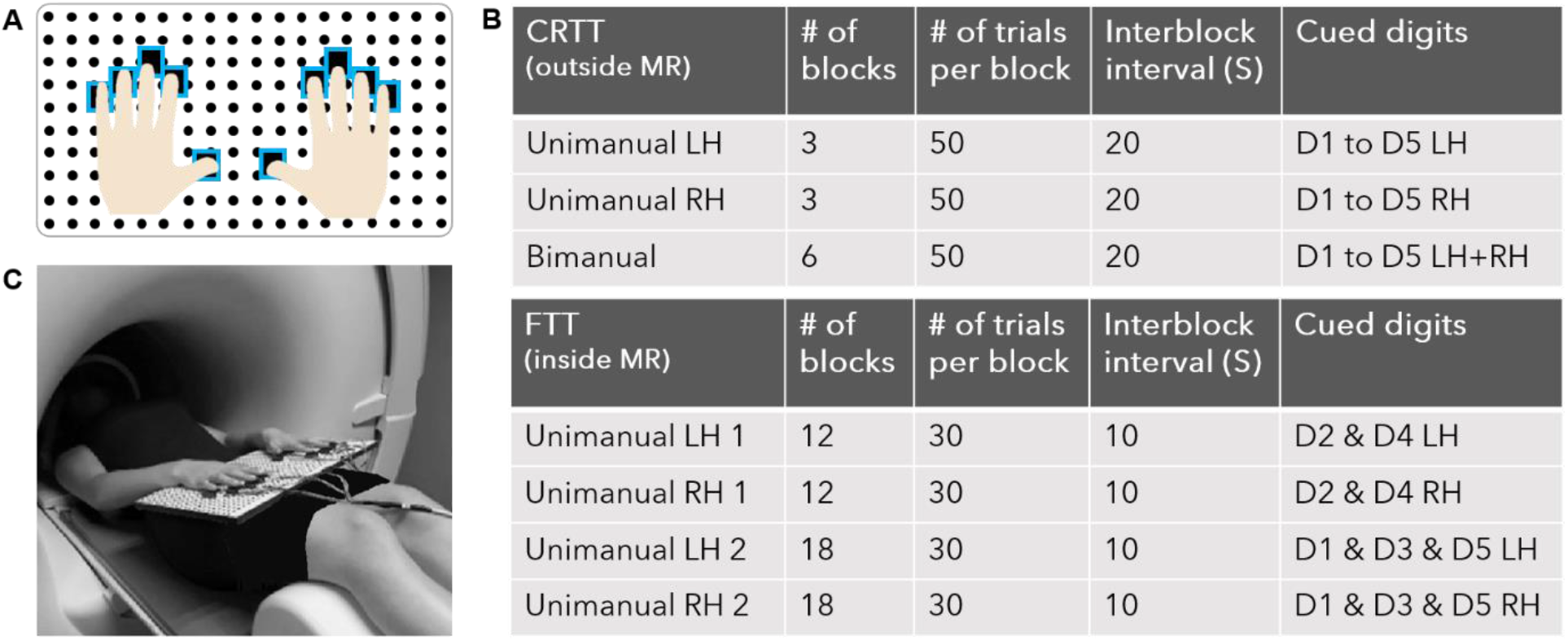
Experimental set-up and task conditions. **Panel A)** The experimental set-up with the fingers positioned on the force sensors. Force sensors were fixated into 3D printed holders with pegs underneath (in blue), which in turn could be placed inside the small holes of the wooden plate. **Panel B)** Overview of all task conditions during the CRTT outside the MR scanner (top) and the FTT inside the MR scanner (bottom). S = seconds; LH = left hand, RH = right hand, D1 = thumb, D2 = index finger, D3 = middle finger, D4 = ring finger, D5 = little finger. **Panel C)** the position of the participant (laying supine in MR scanner) with the experimental set-up resting on the lap of the participant ensuring a comfortable position of the wrists, hands and fingers.

### Finger Tapping Task

In order to assess the relationship between individual digit representations in the sensorimotor cortex and digit enslaving during a simple, continuous movement, we included a Finger Tapping Task (FTT). As this task was also used to localize digit activation patterns in the sensorimotor cortex, the FTT was performed in a supine position on the extendible table of the MRI scanner. The custom-made MR-compatible set-up that was used for the CRTT (see above), was positioned over the participant’s lap at a comfortable distance (Figure 13C). Force transducers were positioned per participant to ensure easy handling of the sensors and a natural position of the fingers and wrists. Small cushions underneath the plate were provided to increase comfort, and foam cushions were placed around the head to limit head movements. The task was displayed via a video projector and viewed via a double mirror, mounted to the head coil. The FTT consisted of four runs according to a block design. Each block started with an instruction screen indicating which finger to move in the upcoming finger tapping block (duration of 2 seconds). Next, a finger tapping block (duration of 20 seconds), and an inter-block rest period (average duration of 10 seconds) followed. During the finger tapping block, a black screen with two white hands and a fixation cross in the center were displayed. The instructed finger for a given trial switched color between white and red at a fixed frequency (1.5 Hz). The participant was instructed to move the corresponding finger [i.e., alternation of extension (fingertip touches the force transducer) and hyperextension (finger is in the air above the force transducer)] according to the visually imposed speed (i.e., 1.5 Hz), resulting in 30 taps of a given finger per finger tapping block. During the inter-block rest period, the visual display was identical, but without the coloring of a finger and participants were instructed to relax their fingers positioned on the force transducers. There were 2 unimanual runs (one for each hand) consisting of 12 finger tapping blocks each, in which tapping of the index finger (digit 2 – D2) was interspersed with the tapping of the ring finger (digit 4 – D4), resulting in 180 taps per finger. In addition, there were 2 runs (one for each hand) consisting of 18 finger tapping blocks each, in which tapping of the thumb (digit 1 – D1), tapping of the middle finger (digit 3 – D3), and tapping of the little finger (digit 5 – D5) were alternated in blocks, resulting in 180 taps per finger. The order of runs, as well as the order of conditions (i.e., digit) within the runs, were pseudo-randomized. An overview of the FTT task conditions is provided in Figure 13B. In order to calibrate the force transducers, baseline measurements (i.e., no fingers on the force transducers) were taken prior to the first, and after the last run. In addition to measuring digit enslaving, the output of the force transducers was used to verify task compliance.

Computer programming for both the CRTT and the FTT was done using National Instruments Laboratory Virtual Instrumentation Engineering Workbench (LabVIEW 2016, 32 bit).

### Imaging procedure

A Siemens Magnetom 7T magnetic resonance scanner with a 32-channel head coil (Nova Medical) was used. High-resolution T1-weighted structural images were acquired using magnetization-prepared 2 rapid acquisition gradient echo (repetition time/echo time = 5000/2.47 ms, voxel size = 0.7 × 0.7 × 0.7 mm^3^, 240 slices; scan time was 9:42 minutes). While participants were performing the finger tapping task, functional images were acquired with a multiband gradient echo (GE) echo planar imaging (EPI) pulse sequence (repetition time/echo time = 2000/18.6 ms, flip angle = 75°, 92 slices, slice thickness = 1.25 mm, in-plane resolution = 1.25 × 1.25 mm, multiband factor 2, GRAPPA acceleration factor 3), phase encoding direction anterior to posterior. In total, the fMRI acquisition took ∼ 35 minutes. In order to correct for off-resonance distortions in the functional images, six additional volumes were acquired with a reversed phase encoding direction (posterior to anterior).

### Analysis

#### Behavioral analysis

The CRTT and FTT data were analyzed using both Matlab (R2018b) and Microsoft Excel (2016). For both tasks, data from all ten force transducers (one for each finger) were analyzed by means of in-house developed Matlab scripts. At first, baseline data (i.e., the signal from force transducers when digits were not on the sensors) were subtracted from the raw signals during task performance, not allowing negative force levels. As baseline data recording of the right thumb for two subjects was problematic, data of the respective digit were removed from further analyses. In addition, one subject did not adhere to the task instructions during the FTT (on which the cortical digit representations are based), resulting in the exclusion of this participant from all further analyses.

### Choice Reaction Time Task

For the CRTT, we obtained the following outcome variables: interference score, digit enslaving, reaction time and accuracy. Each of these is further explained in the section below. Of note, findings on reaction time and accuracy are presented in the Supplementary material.

In order to determine the above mentioned outcome variables, using pwfit, a publically available Matlab script, we fitted a single curve onto the baseline-corrected force data for each digit separately. To detect a correct response, i.e., lifting a cued digit, we searched within the fitted curve for the lowest voltage after presenting the cue. For a correct response, a voltage drop to zero should be present in the fitted curve of the cued digit. Importantly, the CRTT lends itself for measuring accuracy of the cued digits and the non-cued digits at the same time. The former provides an indication of correct excitation, referring to the number of correct trials, i.e., correctly lifting the cued digit (= *accuracy*). The latter provides an indication of the ability to inhibit competing incorrect alternatives. Here, we expressed the latter as an error score (= *interference score*), i.e., counting the number of incorrect lifts of non-cued digits. More specifically, for each non-cued digit we calculated the number of times this digit was lifted incorrectly with the cued digit. The maximum error score is therefore equal to the numbers of trials per digit, i.e., 30. Interference scores were calculated within hand, resulting in a 5 (D1-D5) x 5 (D1-D5) non-symmetrical matrix per hand including 4 interference scores (= all non-cued digits of the same hand) per cued digit, with the diagonal being zero. Moreover, to assess *digit enslaving* in all non-cued fingers relative to the cued finger (within one hand), we followed the procedure reported by Ejaz et al., (2015). During each correct trial (i.e., where the cued finger was lifted from the force transducer), we calculated the sum of the difference in force levels between pre-trial baseline and post-cue presentation for each non-cued finger. Importantly, any change in the continuous force signal of the non-cued digits was included, i.e., digits were not necessarily completely disconnected from the sensor. These co-activation values were arranged into a digit enslaving matrix for the CRTT. Finally, the time elapsing from the cue presentation to a voltage drop in the continuous force signal was calculated as the *reaction time* in milliseconds.

### Finger Tapping Task

For the FTT, we obtained digit enslaving as the sole outcome variable. In order to do so, we analyzed the force data throughout the duration of the task. To find the frequency components of the original force signals (including noise), first a Fast Fourier Transform algorithm was applied. As the imposed movement frequency was 1.5 Hz, we selected all frequency components between 1.0 and 2.0 Hz. Next, signals for each individual digit were smoothed using a Savitzky-Golay finite impulse response smoothing filter. Per trial, for each cued finger, we searched for drops in the smoothed signal, as the digit was lifted from the force sensor. To determine digit enslaving in non-cued digits, correlation coefficients were determined from the sinewaves of each non-cued finger and the sinewave of the cued finger. A positive correlation reflects a co-contraction, i.e., the non-cued finger moves along with the instructed finger, whereas a negative correlation reflects a supporting movement, i.e., the non-cued finger increases the force level onto the transducer to support the continuous tapping of the instructed finger. Correlation values were averaged across blocks of the same cued finger. Correlation coefficients were arranged into a 5 (D1-D5) x 5 (D1-D5) digit enslaving matrix for the FTT.

#### Imaging analysis

Data preprocessing and analyses of functional time series were performed using the BrainVoyager software package v20.6 (Brain Innovation BV, Maastricht, The Netherlands, http://www.brainvoyager.com). To ensure T1 equilibration, the first two volumes from each fMRI run were removed. Pre-processing included slice-timing correction, 3D motion correction to realign all EPIs to the first volume, and temporal high-pass filtering (with a cut-off at 2 cycles, GLM Fourier basis set). Subsequently, EPI distortion correction was performed using the Correction based on Opposite Phase Encoding (COPE) plugin within BrainVoyager (Breman et al., 2020) to correct for geometric distortions. A voxel displacement map was calculated and the image series was corrected with the resulting displacement map. The resulting EPIs were coregistered to the individual anatomical images (gradient-based affine alignment with 9 parameters), resampled to 1.5 mm isotropic resolution, and analyzed by means of a general linear model (GLM). For each experimental run, a design matrix was created including a regressor for each individual digit, for the instruction phase and for the inter-block rest periods, supplemented with six z-transformed motion parameters (covariates of no interest). Per run, blocks were modeled with a boxcar function which was convolved with a hemodynamic response function. Afterwards, all runs were combined in a single-subject multi-run GLM analysis contrasting each individual digit against the baseline. The resulting statistical maps were fed into the representational similarity analysis (RSA) of BrainVoyager, which served to analyze the (dis)similarity of activation patterns evoked by individual finger movements in the sensorimotor cortex using the Pearson correlation method for calculating (dis)similarities. Within each region of interest (see further), a representational dissimilarity matrix (RDM) was computed between pairs of activation patterns representing individual finger movements for each hemisphere and participant separately, as well as across participants. More specifically, for a given brain region, the activation patterns elicited by single finger movements are interpreted as stimulus representations. Next, activation patterns associated by each pair of conditions (i.e., single finger movements) are compared, resulting in an RDM with each cell of the RDM reflecting the dissimilarity (i.e., distance) between the activation patterns elicited by the two single finger movements. As such, the RDM is symmetrical with zeros on the diagonal (Kriegeskorte, Mur, & Bandettini, 2008). The dissimilarity measure *d*, from which the RDM is built, was transformed back to a correlation/similarity value by performing 1 – *d*, such that higher values in the matrix represent more similar spatial activation patterns between single digits. Anatomical regions of interest (ROIs) were defined by means of FreeSurfer (version 21.0, http://surfer.nmr.mgh.harvard.edu). The pial and white-gray matter surfaces of the T1-weighted anatomical image were reconstructed and used to define Broadmann areas (BA) 1, 2, 3a, 3b, 4a, and 4p, on the group surface. The BA 1, 2, 3a, and 3b were then merged to form the primary somatosensory cortex (S1), and BA 4a and 4p were merged to form the primary motor cortex (M1). These ROIs were defined for both hemispheres. As we were interested in individual finger representations at the level of M1 and S1, we selected the hand areas in the pre- and post-central gyri based on a combination of fMRI activation and landmarks (Yousry et al., 1997) and approximately 2 centimeters above and below these regions. Next, M1 and S1 masks were projected onto resampled native space for each participant making use of their individual surfaces (per hemisphere). The resulting ROIs were imported into BrainVoyager (v20.6 – Brain Innovation BV, Maastricht, The Netherlands) to serve as masks to perform the RSA.

#### Statistical analysis

Reaction time data of the CRTT task were analyzed using repeated-measures analysis of variances (ANOVAs) with Hand (left/right), Coordination mode (unimanual/bimanual) and Digit (D1, D2, D3, D4, D5) as factors of interest. As accuracy, interference scores and digit enslaving measures of the CRTT deviated significantly from a normal distribution, non-parametric tests were performed to assess the effects of Hand (left/right), Coordination mode (unimanual/bimanual) and Digit (D1, D2, D3, D4, D5) or Digit pair (vectorized; D1-D2, D1-D3, D1-D4, D1-D5, D2-D3, D2-D4, D2-D5, D3-D4, D3-D5, D4-D5). As matrices including digit enslaving and interference scores were typically not symmetrical, we compared values above and below the diagonal, revealing no significant main effect (*ps* > .05). We, therefore, averaged the values above and below the diagonal and created symmetrized matrices for further analyses. However, as Wilcoxon Matched Pairs Tests comparing digit pairs directly revealed some differences between values above and below the diagonal for interference scores in particular (surviving a Bonferroni correction; *p* < .005), we also performed the analyses for symmetrized data above and below the diagonal separately. Corresponding findings are reported in the Supplementary Materials. In order to compare digit enslaving and interference matrices between the left and right hand, Mantel tests (5000 permutations) were performed for each participant separately, and across participants. The Mantel test is a statistical test of the correlation between two matrices with the same dimensions (Mantel, 1967) and can be applied to determine correlations between matching positions of two (dis)similarity matrices resultant from multivariate data. In addition, to determine the extent to which digit representations in the primary sensorimotor cortices could explain digit enslaving and interference scores in the contralateral hand during the CRTT, subject-by-subject and group neural similarity matrices were compared using Mantel tests (5000 permutations). RDMs of the left hemisphere (M1 and S1) were correlated with enslaving matrices of the right hand, and vice versa. This resulted in four Mantel tests per participant and across participants.

Enslaving data of the FTT (i.e., correlation coefficients) were also analyzed using a repeated-measures ANOVA with Hand (left/right) and Digit pair (vectorized; D1-D2, D1-D3, D1-D4, D1-D5, D2-D3, D2-D4, D2-D5, D3-D4, D3-D5, D4-D5) as factors of interest. As digit enslaving matrices were typically not symmetrical, we compared values above and below the diagonal, revealing no significant main effect (*ps* > .05). We, therefore, averaged the values above and below the diagonal and created symmetrized matrices for further analyses. In order to correlate digit enslaving matrices between the left and right hand, Mantel tests (5000 permutations) were performed for each participant separately, and across participants. In addition, to determine the extent to which digit representational structures in the primary sensorimotor cortices could explain digit enslaving in the contralateral hand during the FTT, subject-by-subject and group similarity matrices were compared using Mantel tests (5000 permutations). As the type of co-activation (i.e., co-contraction (decrease of force on the sensor) or supporting movement (increase of force on the sensor)) is not of relevance when associating with cortical representations, absolute correlation coefficients were used in the following analyses. RDMs of the left hemisphere (M1 and S1) were correlated with enslaving matrices of the right hand, and vice versa. This resulted in four Mantel tests per participant, and across participants. In the results section, correlation coefficients using Spearman’s rank correlation are given along with corresponding *p* values. All Mantel tests were carried out in the R environment using the vegan package (Oksanen, 2012).

## Acknowledgments

This work was supported by the Research Fund KU Leuven (C16/15/070), Fonds Wetenschappelijk Onderzoek Vlaanderen (FWO) (G089818N) and by the FWO-FNRS Excellence of Science Grant (EOS 30446199, MEMODYN). JG and SC are funded by an FWO postdoctoral fellowship. The funders had no role in study design, data collection and analysis, decision to publish, or preparation of the manuscript. The authors would like to thank R. Clerckx for support on the behavioral analysis and J. Diedrichsen and N. Ejaz for support on the non-ferromagnetic experimental setup.

