## Supplementary Material for "Representational similarity scores of digits in the sensorimotor cortex are associated with behavioral performance"

*CRTT outside MR*

*Reaction Time* To describe the performance on the CRTT, repeated-measures ANOVAs with factors Hand (2 levels; left and right), Coordination mode (2 levels; unimanual and bimanual) and Digit (5 levels; D1, D2, D3, D4, D5) were performed. There was no main effect of Hand ($F(1,13) = .19$, $p = .67$), a main effect of Coordination mode ($F(1,13) = 18.40$, $p = .001$), and a main effect of Digit ($F(4,52) = 21.28$, $p < .001$). In addition, the two-way interactions between Coordination mode and Digit ($F(4,52) = 3.06$, $p = .02$), and Hand and Digit ($F(4,52) = 6.86$, $p <.001$) reached significance. Tukey HSD post-hoc tests revealed that for both the uni- and bimanual coordination mode, D1 and D2, D1 and D5, and D3 and D4 revealed comparable reaction times, whereas all other comparisons between digits revealed significant differences, with highest/slower reaction times for D3 and D4. However, whereas for the unimanual coordination mode, D2 and D5 revealed comparable reaction times, for the bimanual coordination mode, D2 revealed higher/slower reaction times than D5. Concerning the Hand X Digit interaction, findings revealed that for the right hand, reaction times for D2 were not significantly lower/faster relative to D3 and D4, whereas this was the case for the left hand. In addition, whereas the right hand did demonstrate faster reaction times for D2 relative to D5, this was not the case for the left hand. Results are presented in Figure S1.

*Accuracy* Regarding median accuracy scores, non-parametric Sign Tests revealed no statistically significant differences between hands ($p = .79$), or between Coordination modes ($p = .45$). There was, however, a statistically significant difference between Digits ($X^2(4) = 25.34$, $p < .001$). Post-hoc Wilcoxon Matched Pairs Tests revealed that D3 demonstrated lower accuracy scores relative to D1, D4 and D5, and that D2 demonstrated lower accuracy scores relative to D5 ($p_s < .05$). Results are presented in Figure S2.

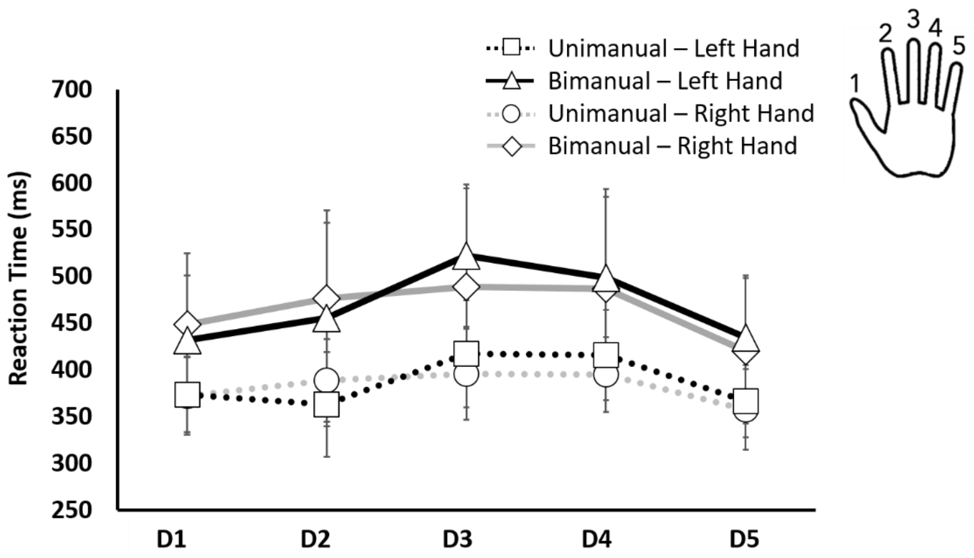


***Figure S1.*** Reaction time scores on CRTT for each digit individually. Error bars indicate standard deviations.

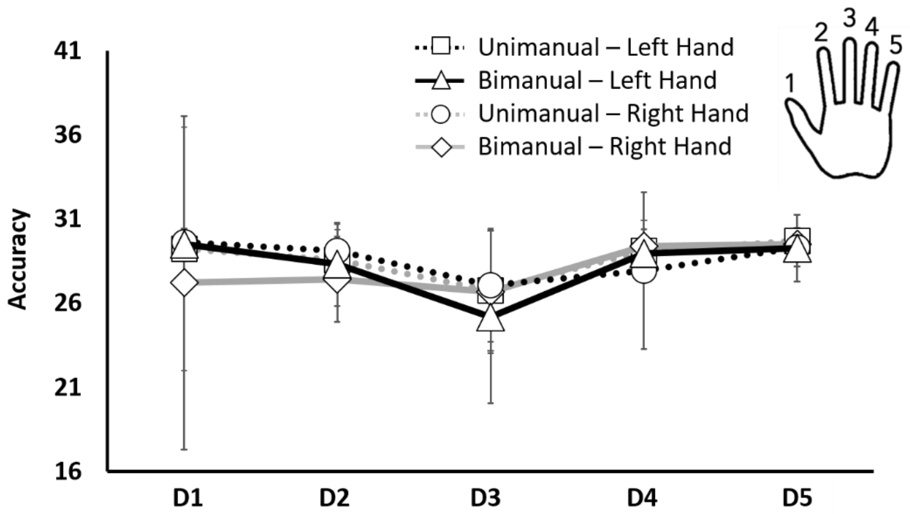


***Figure S2.*** Accuracy scores on CRTT for each digit individually. Error bars indicate standard deviation.

*Digit interference scores above and below diagonal associated with representational structures*

As Wilcoxon Matched Pairs Tests revealed significant differences in CRTT interference scores between values above and below the diagonal (i.e., which digit of the respective digit pair was cued) for digit pairs D1-D2, D2-D3, D2-D4 and D2-D5, we decided to study the association with neural representations for the values above the diagonal (symmetrized) and for the values below the diagonal (symmetrized) separately. Mantel tests with 5000 permutations revealed significant positive associations between left hand interference scores above the diagonal and contralateral S1 ($r$ = .54, $p$ = .03) and M1 ($r$ = .56, $p$ = .03). In addition, right hand interference scores above the diagonal were significantly associated with contralateral S1 ($r$ = .67, $p$ = .04), and a trend for M1 ($r$ = .65, $p$ = .06). Associations concerning interference scores below the diagonal did not reach a statistical significance level ($p_s > .05$). Since the digit pairs demonstrating a significant difference between values above and below the diagonal all include the index finger (D2), the origin of differential associations with neural representations is likely to be found there. Values above the diagonal refer to digit pairs in which the first digit is cued. Since the index finger is known to behave rather independently from other fingers within the same hand, it is understandable that the index finger is unlikely to co-move when other fingers are cued. At the same time, it is likely that less independent fingers such as the middle and ring finger do show co-contractions when the index finger is cued. The fact that significant associations are mainly found in cases where the index finger is cued, and neighboring fingers are co-contracting, likely reflects that this phenomenon better resembles the digit representations at the contralateral cortical level.
